# Temperature-dependent Self assembly of biofilaments during red blood cell sickling

**DOI:** 10.1101/2021.03.07.433773

**Authors:** Arabinda Behera, Oshin Sharma, Debjani Paul, Anirban Sain

## Abstract

Molecular self-assembly plays vital role in various biological functions. However, when aberrant molecules self-assemble to form large aggregates, it can give rise to various diseases. For example, the sickle cell disease and Alzheimer’s disease are caused by self-assembled hemoglobin fibers and amyloid plaques, respectively. Here we study the assembly kinetics of such fibers using kinetic Monte-Carlo simulation. We focus on the initial lag time of these highly stochastic processes, during which self-assembly is very slow. The lag time distributions turn out to be similar for two very different regimes of polymerization, namely, a) when polymerization is slow and depolymerization is fast, and b) the opposite case, when polymerization is fast and depolymerization is slow. Using temperature dependent on- and off-rates for hemoglobin fiber growth, reported in recent in-vitro experiments, we show that the mean lag time can exhibit non-monotonic behaviour with respect to change of temperature.

## INTRODUCTION

Self-assembly of different types of protein filaments is essential for the structural organization and functioning of the cell. Important examples are actin and microtubule in eukaryotes and FtsZ filaments in bacteria. However, abnormal self-assembly or aggregation of certain proteins are known to cause various diseases such as Parkinson’s disease, Huntington’s disease, prion diseases, Sickle cell disease (SCD) [1–3] etc. Abnormal self-assembly typically leads to the formation of stiff protein fibers, fiber bundles, and branched structures that can deform the cell. For example, red blood cells deform into a sickle-like shape [4, 5] in SCD when abnormal hemoglobin (HbS) proteins, in an oxygen-deficient environment, polymerize to form long rigid fibers. The sickled RBC loses its elasticity and fails to pass through the narrow capillaries, resulting in vaso-occlusion. Although we will primarily focus on the kinetics of self-assembly of HbS proteins, some of our findings may be universal across other fibrillating proteins.

A significant delay or a lag time in the formation of the polymerized mass has been one of the widely studied features of this kinetics. Longer lag time is beneficial for SCD as it allows the deoxygenated RBC to return from the capillary before it sickles. In vitro experiments monitor the growth of polymerized protein mass Δ(*t*) after artificially triggering the fibrillation at time *t* = 0. Beyond the initial lag time, Δ(*t*) shoots up exponentially and saturates after that. This delay, of the order of few minutes, has been identified as the nucleation time of HbS aggregates/oligomers, which thereafter grows/polymerizes rapidly at the rate of a few *μm* per seconds [6] and deform the RBC. Interestingly, the lag time shows wide stochastic variation. Both the mean lag time and its stochasticity show systematic dependence on monomer concentration and temperature [7], which is the focus of our simulation study here.

Stochasticity has been attributed to small reaction volume and low HbS concentration of the fibrillating proteins. Classic experiments with RBC [8], and recent in vitro microdroplet experiments on bovine insulin fibrillation [9, 10], also support this point. Ref.[10] suggested that lag-time varies inversely with the volume of the system. In fact, Ref[11] has suggested that artificial volume expansion of RBC (causing a decrease in the HbS concentration) could be an effective therapeutic tool for extending lag time.

Nucleation events are always stochastic; it is a thermally activated barrier crossing process. How does reaction volume affect the stochasticity of the lag time? Small reaction volume has a proportionately small number of available monomers as well as nucleation events. Once the first nucleus forms, its rapid growth/polymerization depletes monomers in the medium, making further nucleation events difficult. On the other hand, in a large reaction volume, since the monomer pool is large, multiple nuclei can form at different spatial points and grow, making the lag time less stochastic.

The main focus of this article is the dependence of the lag time on temperature and monomer concentration. Early experiments [6, 12] have highlighted this and estimated how monomer association *k_on_* and dissociation *k_off_* rates may vary with temperature, influencing the lag time. A recent experiment by Castle et al. [13] have claimed that these kinetic coefficients had been critically underestimated before (by order of magnitude). The temperature dependence they found is quite counterintuitive: *k_on_* increases with temperature while *k_off_* decreases with temperature. Naively one would expect that *k_off_* must increase with temperature as breaking a bond is easier at higher temperatures. Here we show that if these rates are assumed to be valid during the nucleation process, then the lag time can show non-monotonic behavior with increasing temperature. It is well known that both lag time and its stochasticity increases drastically at low (< 20°*C*) temperatures [12]. However, there are no reliable data at higher than room temperature [11].

Two new aspects of the HbS fibrillation process dictate the choice of our model here. a) Recent experiments on both amyloid fibrillation and RBC sickling showed that nucleation is always preceded by the formation of a dense liquid-gel-like droplet from the monomers [14, 15], beyond a specific concentration. These dense droplets are known to promote protein fibrillation [14–16], which naturally enforces small reaction volume (size of the droplets) and high local concentration of monomers. Therefore, models which ignore the spatial extent of the system (i.e., role of diffusion) and assume that all reactions take place at one point, are reasonable choice for quantitative description.

b) The 2nd point is that most theories [10, 12, 17, 18] assume a given rate (*α*_1_) for primary nucleation and a critical nucleus size *n_c_* = 2. Since lag time is defined as the time taken to build up a certain fraction of the final polymerized mass (eg, 1*/*10), its stochasticity mainly arises from the stochasticity of the early growth, secondary nucleation via fragmentation of the fibrils and their subsequent autocatalytic growth [10, 12, 17, 18]. In contrast, experimental measurements by Galkin et al [6] estimate, for HbS in RBC, *n_c_* ≃ 11 − 12. For such a large *n_c_* we cannot assume the process to be an one-step assembly reaction leading to a nucleation rate 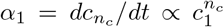 where *c*_1_ and 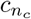 are the concentrations of monomers and nucleus, respectively. If nucleation occurs by a sequential polymerization process as described in Ref[4] this could contribute large stochasticity to the lag time. This is one of the questions we explore here using kinetic Monte-Carlo (KMC) simulations. We take the KMC approach used by Yuvinec et al. [19, 20] who studied nucleation time, defined as the time it takes to form a *critical nucleus*, a cluster of specific size *n_c_*, for the first time, starting at time *t* = 0 with a state which has only monomers.

The paper is organized in the following way. In section-2, we discuss experimental data on the statistics of RBC sickling time, mostly from the literature and some of our data from in vitro studies of RBC sickling. In section-3, we introduce our simulation model for nucleation. Since much of our focus is on the mean lag time from our stochastic simulations, in section-4, we compare data from our stochastic model with the results from the deterministic Becker-Döring (BD) model. Section-5 describes our Kinetic Monte Carlo (KMC) scheme. Our main results are discussed in section-6. We conclude with a discussion section.

## EXPERIMENTAL DATA

In fig1, we show our experimental data for the sickling time (*τ_s_*) measured for 526 RBCs, collected from four different sickle cell disease patients, at room temperature. The experimental conditions have been described in detail in ref[21]. Dashed vertical lines separate these four sets of data on the x-axis. These patients’ blood differs in their HbF, HbA0, HbA2, and HbS content; their sickling time variance was also different. However, as shown in the figure, the most frequently occurring sickling time (mode) was close in all the patients except the 1st one. In order to plot the distribution of *P* (*τ_s_*), shown in fig1-a, we shifted the data of the 1st patient up by 0.35 min to bring its mode at the same level as the other patients. *P* (*τ_s_*) shows a tail although we have truncated the distribution for large *τ_s_* due to lack of sufficient data. In our experiment, the chemical oxygen scavenger (0.1% sodium metabisulphite) was added to these four blood samples at time *t* = 0, and the subsequent time evolution of a large number of RBCs in the sample was recorded at 30 fps under a microscope. The sickling time was defined as the onset of RBC deformation, which was detected visually. After the onset, it took about 5 seconds for the full deformation to complete [5]. The uncertainty in visually detecting the onset was therefore small compared to the typical deformation time, which ranges from 1/2 a minute to 4 minutes. We assumed that the onset of RBC deformation takes approximately the same time as the nucleation since exponentially fast polymerization occurs post nucleation, as mentioned earlier. The deformation occurs when the growing HbS polymers hit the RBC surface. Similar sickling times were reported for in vitro experiments, at room temperatures, and at a physiological concentration of HbS [4, 8, 22].

**FIG. 1.**
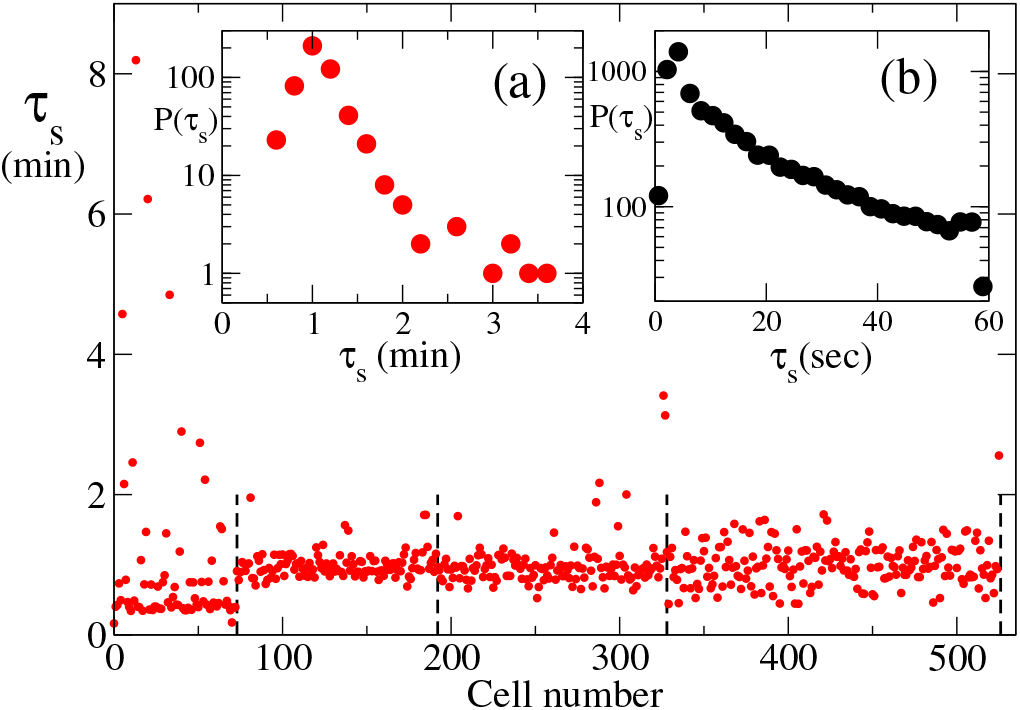
Measured sickling times for about 550 RBC collected from four patients. (a) shows the corresponding distribution, and (b) shows the distribution from a much bigger study from Ref[11] where sickling was induced by laser pulse.

In fig1inset-b we show a distribution obtained in ref[11] using a much larger data set. However, in this in vitro experiment, sickling was induced via laser polymerization in sickle trait RBCs, which leads to much more rapid sickling (note the time scales for insets-a and b). This benchmark distribution shows an exponential behavior. In this experiment, the sickling time was determined by quantitatively measuring three geometric markers of RBC deformation as functions of time.

## OUR STOCHASTIC MODEL AND SIMULATION SCHEME

Figure2 shows the schematics of our nucleation model. Starting from a population of free monomers at *t* = 0, the system goes through the polymerization process to form intermediate aggregates. The simulation is stopped when an oligomer of a predefined size (the nucleus) forms for the 1st time and the corresponding time is recorded as the nucleation time *τ_s_*.

**FIG. 2.**
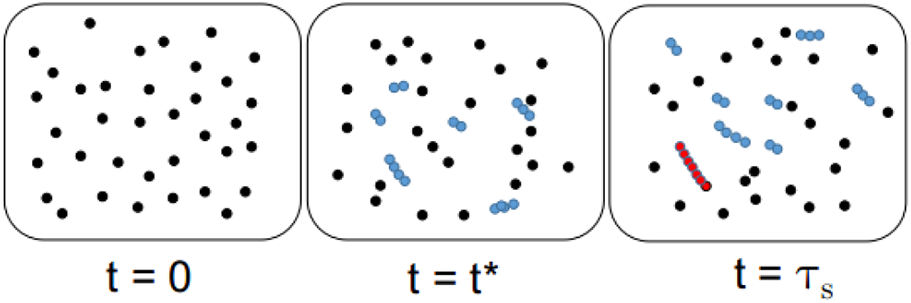
A schematic for the nucleation time for the formation of critical nucleus. The first panel shows the initial state (*t* = 0) where there are only monomers. The middle panel is the intermediate state (0 < *t*^∗^ < *τ_s_*), where there are filaments of different sizes, but no nucleus. The third panel is the final state (*t* = *τ_s_*) where a single nucleus of size 7 has formed. For nucleus with multiple filaments we record the time when multiple disjoint filaments cross the threshold length.

The nucleation events in a small volume (≈ 10^−10^*cm*^3^) is stochastic in nature[12], which is the reason behind the broad sickling-time distributions observed in experiments (see fig.1, and ref [12, 22]). We use Gillespie algorithm[23] to simulate the homogeneous nucleation. Our model is similar to the models discussed in ref.[19, 20]. As observed earlier[5], the HbS polymers grow(shrink) by association(dissociation) of monomers and not oligomers. So, in our model, we consider a single association/dissociation of monomers at a time. We consider the following reactions to the formation of the critical nucleus.

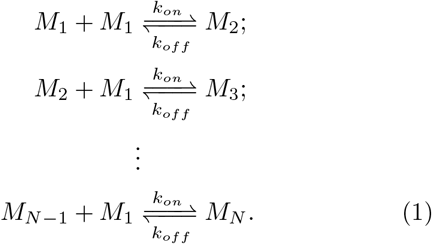

Where, *M*_1_ is the number of monomers, *M*_2_ is the number of dimers and so on. N is the nucleus size which we fix in each simulation. *k_on_* is the on-rate and *k_off_* is the off-rate of the polymers. The simulation stops when the first nucleus forms. So, *k_off_* = 0 for last reaction i.e., there is no detachment from *M_N_*.

For the simulation of eq.1, we use the Gillespie algorithm[23]. First, we define propensity(*α*) for each of the 2*N* − 2 reactions. The propensity of each reaction is proportional to the number of reactants. Then the total propensities of the reactions is given by, 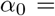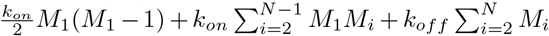. The time step at which the next reaction will occur is drawn from a distribution, 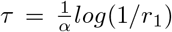, where *r*_1_ ∈ (0, 1) is a computer-generated random number. Then for the *N ^th^* reaction to occur, we draw another random number *r*_2_ ∈ (0, 1) which has to satisfy 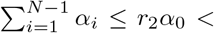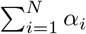, where *α_i_*’s are the propensities of each reaction. Fig.2 shows the schematics of our model. Starting from all monomers at *t* = 0, the system forms the nucleus at *t* = *τ_s_*. The model discussed in ref.[19] has studied the off-rate dependencies in the distribution of nucleation time, keeping the on-rate fixed. We use the same model, but we vary both on-rate and off-rate as a function of temperature. Our model was motivated by the recent experimental study[13], where the authors observed that the on-rate and off-rate depend on the initial monomer concentration and temperature. Also, we will study the case when multiple nuclei form. This could be a proxy for forming a single nucleus with multiple strands, provided the multiple strands in a single nucleus are assumed to grow/shrink independently of each other.

Before presenting our results on the distributions (fluctuations) of nucleation times, from our stochastic simulations, it is useful to know how the mean nucleation time behave. For this, we will compare our mean observables with the predictions of the standard Becker-Döring theory of homogeneous nucleation.

## MEAN NUCLEATION TIME : BECKER-DÖRING MODEL

When adapted for nucleation, the BD equations are a set of differential equations describing the population dynamics of clusters/polymers of all sizes, up to the nucleus size. When monomer addition or subtraction is the only allowed processes responsible for growth or shrinkage of the clusters, the relevant equations are [19, 24],

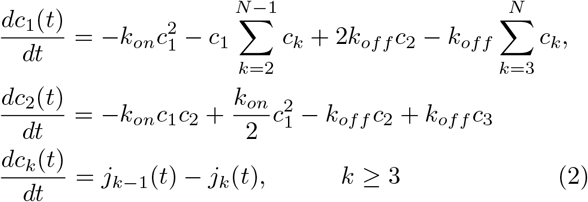

where, *j_k_*(*t*) = *k_on_c*_1_(*t*)*c_k_*(*t*) − *k_off_ c_k_*_+1_(*t*). Here, *c_k_* is the number of clusters of size k, and *c*_1_ is the number of monomers. Here, *k_on_* and *k_off_* are on-rate and off-rate respectively. For simulation of critical nucleus, we start with monomers and set other polymer population to zero, i.e., *c*_1_(0) = 1000 and *c_k_*(0) = 0 for *k* ≥ 2. Here, for simplicity we set the on and off rates, as well as the volume to 1. This makes *c_k_* the number density. As we consider the growth of polymers upto critical nucleus size, we restrict the growth at critical nucleus size N i.e. the simulation stops when *c_N_* (*t*) = 1. The simulation results for eq.2 is given in fig.3.

**FIG. 3.**
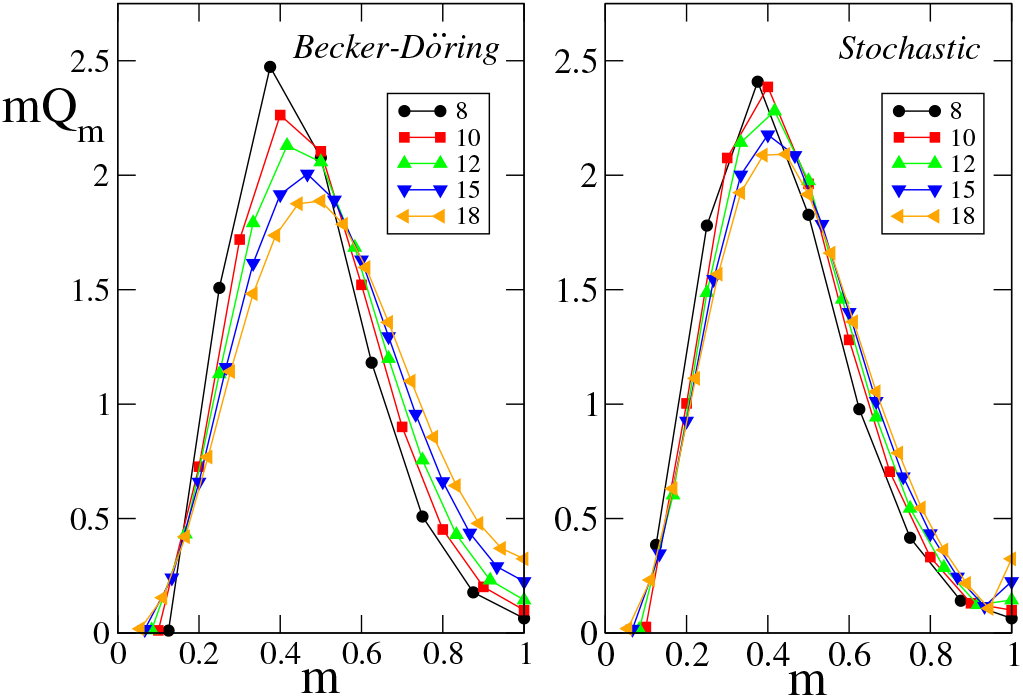
Distribution of filament lengths *m* = *k/N* when the first nucleus forms. The distribution *Q_m_* = *N* ^2^*c_k_/N_T_* is suitably scaled by nucleus size *N* and total number of monomers in the system *N_T_*, such that *mQ_m_* is normalised. Data for five different nucleus lengths are shown, for BD model in (a), and from KMC simulations in (b). Results from both the models are similar, but the KMC results show a better data collapse.

Figure 3a shows the average mass distribution in various sized clusters when the 1st nucleation occurs. The total number of monomers stored in a cluster of size *k* is *kc_k_*, while the nucleus is defined as a cluster of size *N*. The simulation is stopped when *c_N_* reaches 1 for the 1st time. Note that *c_k_* is a continuous variable in Becker-Döring (BD) model, while in the stochastic simulations *c_k_* are integers. *N* was varied for different runs (see legends in 3). In stochastic simulations, for each *N*, the data was averaged over 10^4^ runs. Both BD equations and stochastic simulation conserve mass, i.e., the total number of monomers involved. For all runs, which may differ in nucleus size *N* (see the legends), the systems had the same total mass *N_T_* = 1000. This is achieved by setting the number of monomers at the beginning of the simulation *c*_1_(*t* = 0) = *N_T_*. From mass conservation, we have 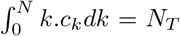. To represent the data from runs with different nucleus size *N* on the same footing, we define rescaled variables *m* = *k/N* and *Q_m_* = *N* ^2^*c_k_/N_T_*, such that 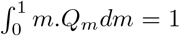, i.e., the area under all the curves are same, irrespective of the nucleus size *N*. Although at nucleation *c_N_* = 1 for all the cases, due to rescaling, the corresponding *Q_m_*_=1_ = *N* ^2^*/N_T_*, differs with *N*. Note that due to rescaling and averaging over many runs, *m* and *Q_m_* both are now continuous variables. The data differing in nucleus size shows a reasonable collapse. The stochastic simulation shows a better data collapse compared to the results from the BD model. Interestingly both simulations reveal that, at nucleation, the maximum number of monomers are engaged in clusters of size about 0.4*N*. We do not have any theoretical explanation for this finding.

Being a deterministic model the BD equations cannot describe the sickling time distribution observed in experiments (see fig.1). But so far as the mean nucleation time ⟨*τ_s_*⟩ is concerned, results from the BD model match with those from the stochastic model. An example is shown in fig.4-a. We found similar match between BD and our stochastic models for the results in fig.4-b as well. So BD model can be conveniently used to study how ⟨*τ_s_*⟩ would change if the on and off rates (*k_on_* and *k_off_*) are made to vary with temperature [13]. Our results in fig.4 shows that, ⟨*τ_s_*⟩ depends separately on both on and off rates, *k_on_* and *k_off_*. Fig.4-b shows a non-monotonic behavior in ⟨*τ_s_*⟩ with respect to varying *k_off_*, with *k_on_* fixed. We varied the ratio in two different ways, a) by varying both *k_on_* and *k_off_*, but keeping their ratio fixed, and b) by increasing *k_off_*, keeping *k_on_* fixed, i.e., increasing the ratio. The results differ for the two cases, and we find non-monotonic behavior for the latter case.

**FIG. 4.**
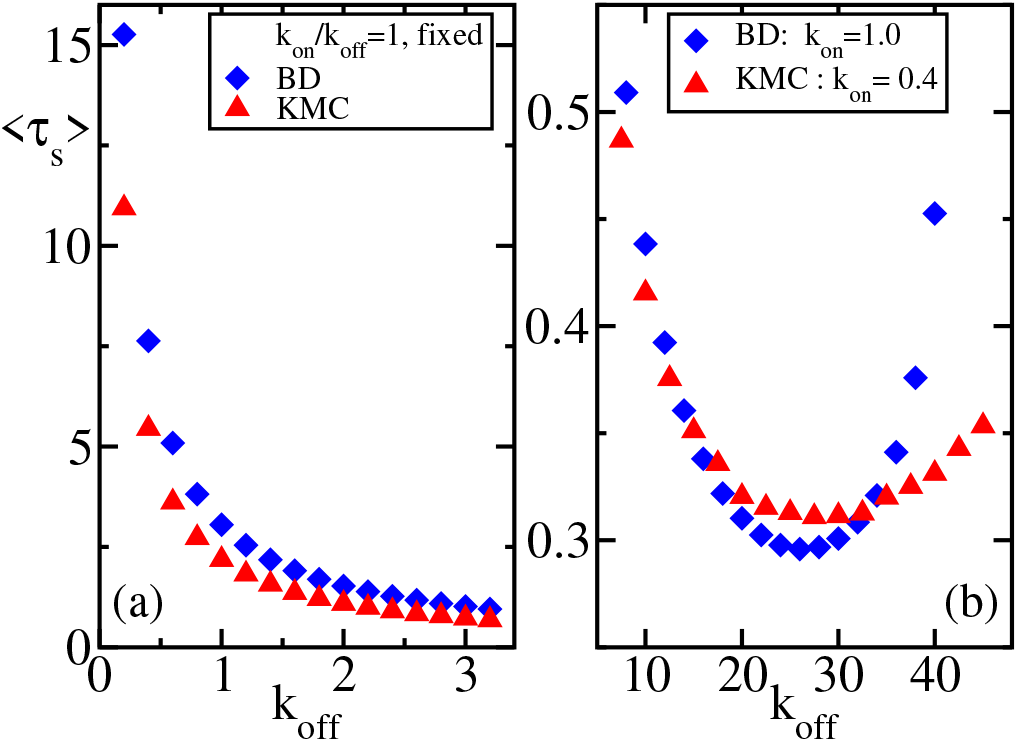
Mean nucleation time ⟨*τ_s_*⟩ as a function of *k_off_*. (a) Shows that ⟨*τ_s_*⟩ decrease monotonically if both *k_on_, k_off_* are increased. This is because a jammed state due to rapid polymerization is avoided by increasing *k_off_*. Note that since the two rates do not have the same dimension, the effective polymerization slows down due to depletion of monomer concentration as time progresses. (b) Non-monotonic dependence of ⟨*τ_s_*⟩ at relatively high *k_off_*. High ⟨*τ_s_*⟩ on the left side of the minima correspond to jammed states when *k_off_* is relatively low, whereas the right side corresponds to relatively slow polymerization (i.e., smaller *k_on_/k_off_*). Our distributions of *P* (*τ_s_*), later, will mostly focus on the left side, i.e., on jammed states.

This non-monotonic dependence has been reported before [19]: at low-*k_off_*, the system quickly gets into a jammed state when all the monomers are used up in forming sub-critical oligomers (of size *k* < *N*). Further growth of these oligomers, towards critical size, depends on effective monomer exchange among these oligomers, i.e., when a monomer released by one oligomer is captured by another oligomer. Because of low off-rate (i.e., slow release) this process slows down. On the other hand, at relatively high off-rate, the sub-critical oligomers are highly unstable and cannot grow further easily.

Going beyond ⟨*τ_s_*⟩ we now discuss the temporal fluctuations using our stochastic KMC simulation. As mentioned earlier [12], in vitro sicking in small volumes are found to be stochastic. Since these volumes are comparable to a actual RBC volume, stochasticity in *τ_s_* is expected inside RBC also.

## SIMULATION RESULTS

### Sickling time distribution

Although the nucleation model which we study here [19] is different from the generic theoretical models of fiber nucleation and growth [10], the lag time distributions of these models turn out to be very similar. In the generic models large stochasticity of lag times arise when, a) number of nuclei are small, and b) they are sparsely spaced in time. The model pioneered by Szabo [17] yield the following distribution for lag times

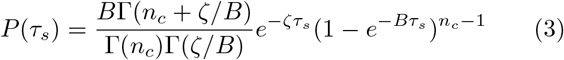

Here *ζ* is the nucleation rate, *n_c_* is proportional to the total polymerized mass at saturation (in fact 1/10-th of that) and *B* is the net rate of mass addition via polymerization/dissociation/splitting of oligomers. This expression has successfully fitted experimental data, which show a rapid rise at small *τ_s_* and a long exponential tail at large *τ_s_*. The time scale *ζ*^−1^ is responsible for the tail, while *B*^−1^ and *n_c_* are responsible for the rapid initial rise. The lag time distributions *P* (*τ_s_*) from our model fits very well with this expression, as shown in fig.5, provided we interpret *n_c_* to be the nucleus size *N* in our model. As *n_c_* increases *P* (*τ_s_*) becomes more skewed and develops long exponential tails (see fig.5c). As in Szabo’s model [17], our *P* (*τ_s_*) is more skewed for larger nucleus size (*n_c_*) and yields larger values of the time scales *B*^−1^ and *ζ*^−1^, which are fit parameters. An increase of both these time scales increases both the standard deviation (*σ_N_*) and the most probable values 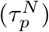 of the lag times. In the inset of fig.6, we show these features. In fact *σ_N_* shows a clear exponential rise with nucleus size *N*. In fig.6, we show the normalized and rescaled *P* (*τ_s_*) for different nucleus size *N*. Maxima of the distributions, for the different nucleus sizes (*N*) are shifted to zero and the spreads around the maximum are rescaled by respective *σ_N_*, by using the variable (*τ_s_* − *τ_N_*)*/σ_N_*. A moderate collapse is obtained this way, although it is evident that *P* (*τ_s_*) is more pointed near the peak at larger *N_c_*.

**FIG. 5.**
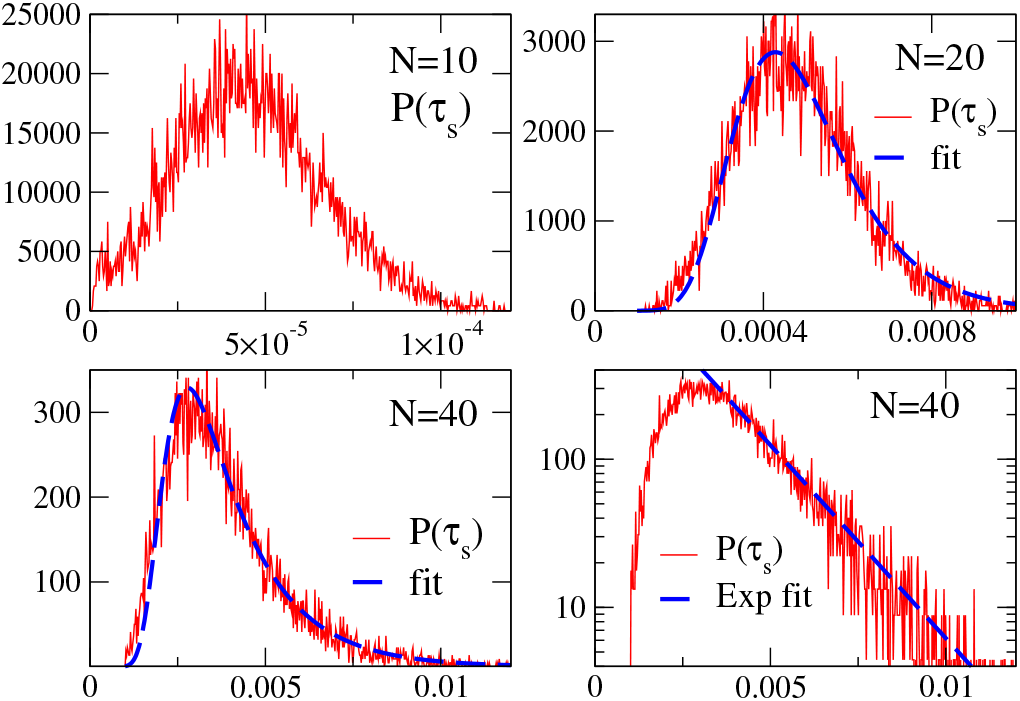
The distribution *P* (*τ_s_*) of nucleation times *τ_s_*, for various nucleus sizes *N* (see legends). For each case initial number of monomers was *M* = 500, with *k_on_* = 4600 and *k_off_* = 35000. Each distribution was obtained from 10^4^ nucleation events. In the lower panels the same data was plotted using regular and semilog scale to highlight the exponential tails. *P* (*τ_s_*) grew more skewed for larger *N* and were fitted to Szabo’s formula Eq.3. With *n_c_* = *N*, both the fit parameters *ζ*^−1^ and *B*^−1^ got smaller for larger *N*.

**FIG. 6.**
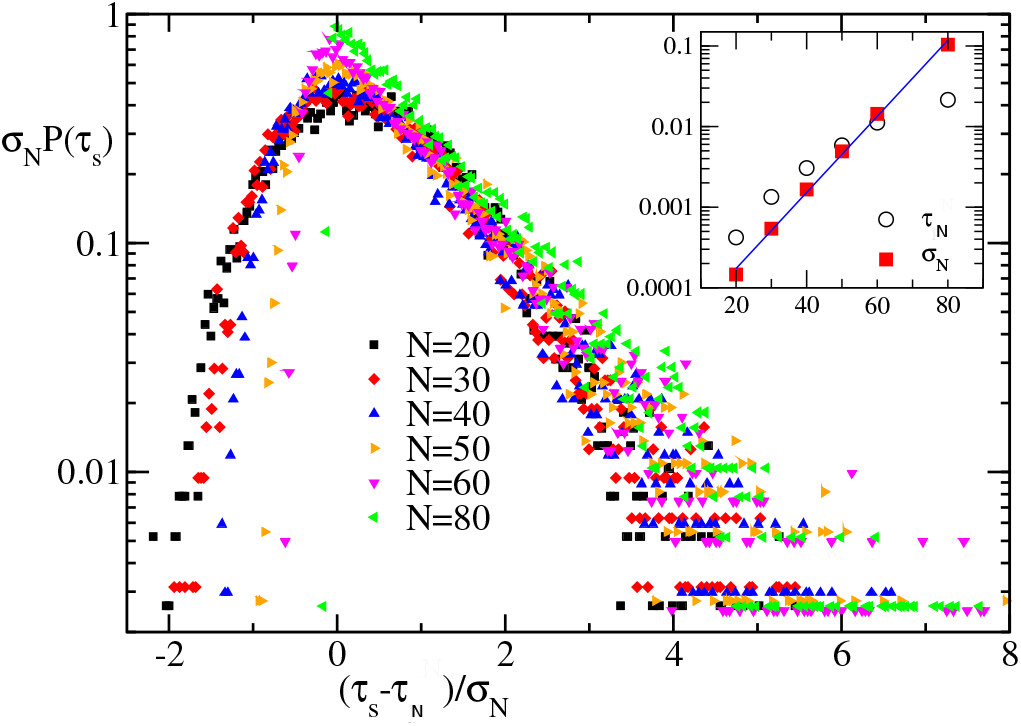
Distribution of nucleation times for various nucleus sizes *N* (see legends). An approximate data collapse is attempted by shifting the peak positions *τ_N_* to zero and rescaling the deviations from the peak positions by *σ_N_* and then maintaining normalization by rescaling *P* (*τ_s_*) by *σ_N_*. At higher *N* (≥ 50), *P* (*τ_s_*) is more pointed near the peak. The inset shows that *σ_N_* scales exponentially with *N*. These distributions correspond to the jammed states where *k_on_* is high and *k_off_* is low.

The apparent similarity between these models, we believe, result from the long waiting times encountered at large *N*. In our model, it is due to the depletion in the number of available monomers, which slows down the growth of the long polymers leading to a trapped state. Whereas in Szabo’s model [17], the low nucleation rate *ζ* and low polymerization rate *B* (at low monomer concentration) causes large waiting times. Even in our simulation model, larger the *N* larger the depletion of monomer concentration, leading to the lowering of *B*.

### Concentration dependence

Our simulations show that high initial monomer concentration can significantly lower the waiting times. That occurs when monomers concentration is not significantly depleted by the time the threshold length of *N* is reached. We show this effect in fig.8 for fixed *N* = 30 and for different initial monomer concentrations (*M*). A near collapse of *P* (*τ_s_*) is achieved for higher values of *M*. The exponential tail is visible only for the lowest *M* value. Interestingly, the skewness shows a change of sign as *M* is varied.

**FIG. 7.**
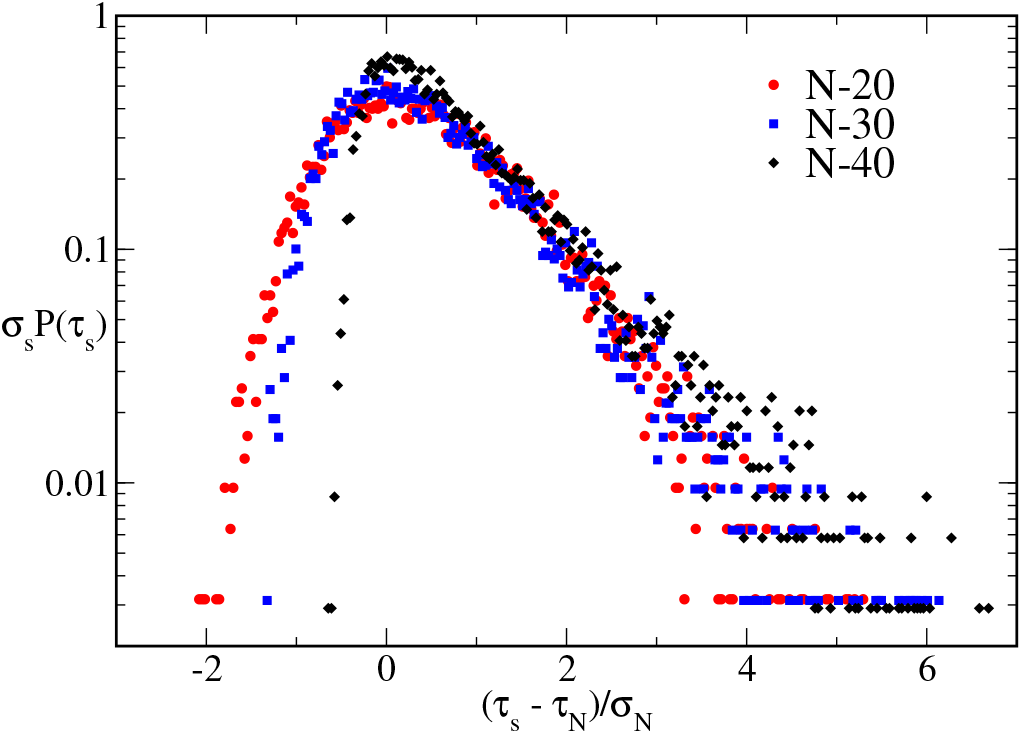
Distributions *P* (*τ_s_*), similar to fig.6, but corresponds to the slow polymerization regime, i.e., low *k_on_* and high *k_off_*.

**FIG. 8.**
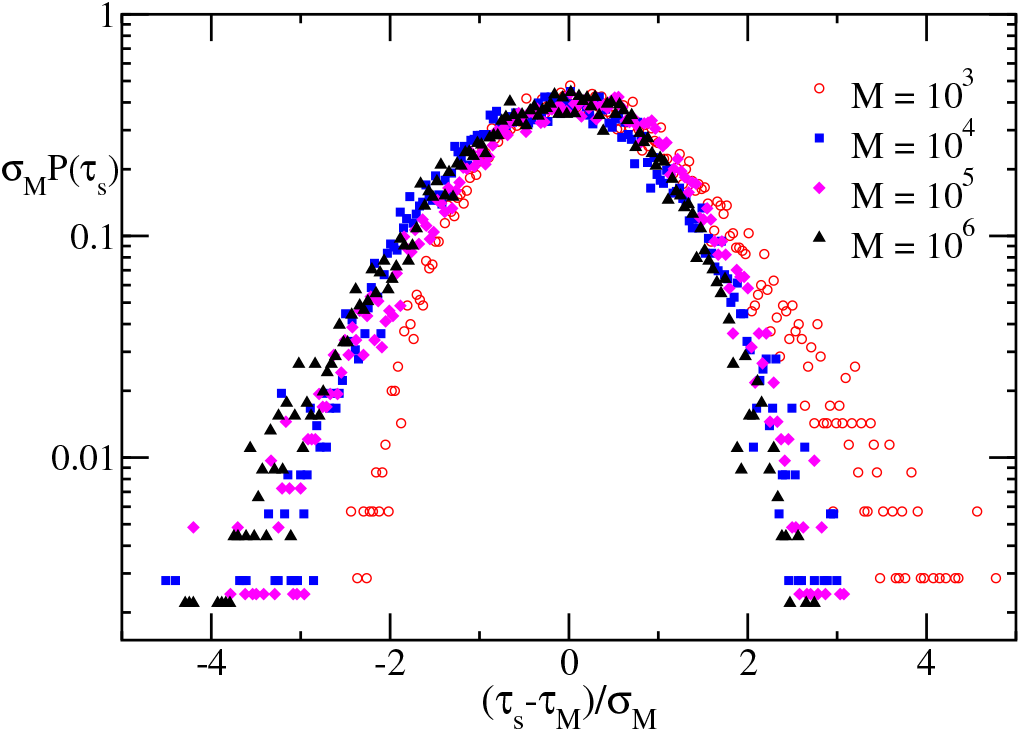
Distributions *P* (*τ_s_*) for different initial monomer populations *M*, and for fixed *N* = 30. The parameters correspond to the jammed state regime where *k_on_* is high and *k_off_* is low. At higher *M*, the skewness changes sign. At lower *M*, the PDFs are similar to those in the jammed regime.

In fig.9, we show another result from our model, which is qualitatively similar to the experimental results. The average lag time ⟨*τ_s_*⟩ in experiments[8] vary over few orders of magnitudes when the initial monomer concentration is varied by 20%. In our model, the ⟨*τ_s_*⟩ reduces by 20 times due to a ten-fold increase in the monomer concentration. Also, the standard deviation of the distributions increases 2.6 times with a decrease in monomer concentration by ten times. A qualitatively similar trend was observed in ref.[25].

**FIG. 9.**
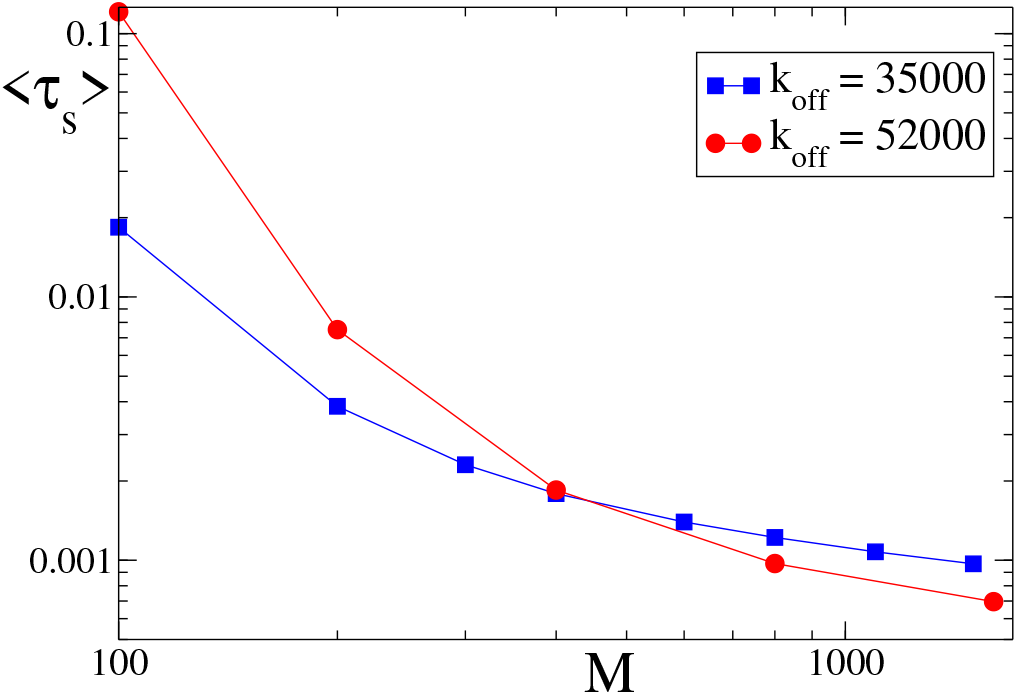
Rapid variation of mean sickling time ⟨*τ_s_*⟩ as a function of initial monomer population *M*, for nucleus size *N* = 30. The two curves, with squares and circles, belong to the low off-rate and high off-rate regimes, respectively, i.e., they correspond to the jammed state and slow-polymerization state. The solid lines are a guide to the eye. Our results are qualitatively similar to the experimental observations in ref[4], where the *t*_1/10_(similar to lag time) decreases by six orders of magnitude due to a 2-fold increase in monomer concentration.

### Bundle formation

We also considered the case of HbS bundle formation during nucleation. Cryo-em studies [26] have confirmed the existence of HbS bundles which form via inhomogeneous nucleation. In inhomogeneous nucleation, an existing oligomer’s surface serves as the nucleation site for more oligomers, which grow parallel to the original oligomer and eventually form a bundle. We mimicked this situation by stopping the simulation when multiple polymers of specific length *N* has formed. For example, when three polymers cross this threshold length *N* (not necessarily simultaneously), we call it a nucleus. Since we do not have the notion of space in our simulation, it does not matter whether these three polymers are growing side by side or not. However, if there is any cooperative effect in bundle formation, our simulation does not capture that. As expected, we find that the lag time gets extended, and *P* (*τ_s_*) becomes more skewed with a longer exponential tail (figure not shown here). This is because when the 3rd longest polymer reaches length *N*, the longest polymer has reached a length *N*′ > *N*, which is equivalent to the lag time of a mono-filament nucleus of size *N*′.

### Temperature dependency

As reported earlier[13], the monomer on- and off-rates are both temperature-dependent. As the temperature is increased, the on rate increases, and the off-rate decreases. As a result, how does the mean nucleation time (*τ_s_*) respond to increasing temperature? Equivalently, within our KMC simulation, we study how *τ_s_* respond to change of on-rate (and corresponding off-rate).

Figure10 shows the variation of ⟨*τ_s_*⟩ with the on-rate. In ref[13], it has been observed that the on-rate varies from *k_on_* = 3000*μM* ^−1^*s*^−1^ to *k_on_* = 4600*μM* ^−1^*s*^−1^, as the temperature increases from 25°*C* to 35°*C* and accordingly the off-rate decreases from *k_off_* = 52000*s*^−1^ to *k_off_* = 35000*s*^−1^. The units in our KMC simulation are not same as in the real experiment because in our simulation all the reactions occur essentially at the same point.

**FIG. 10.**
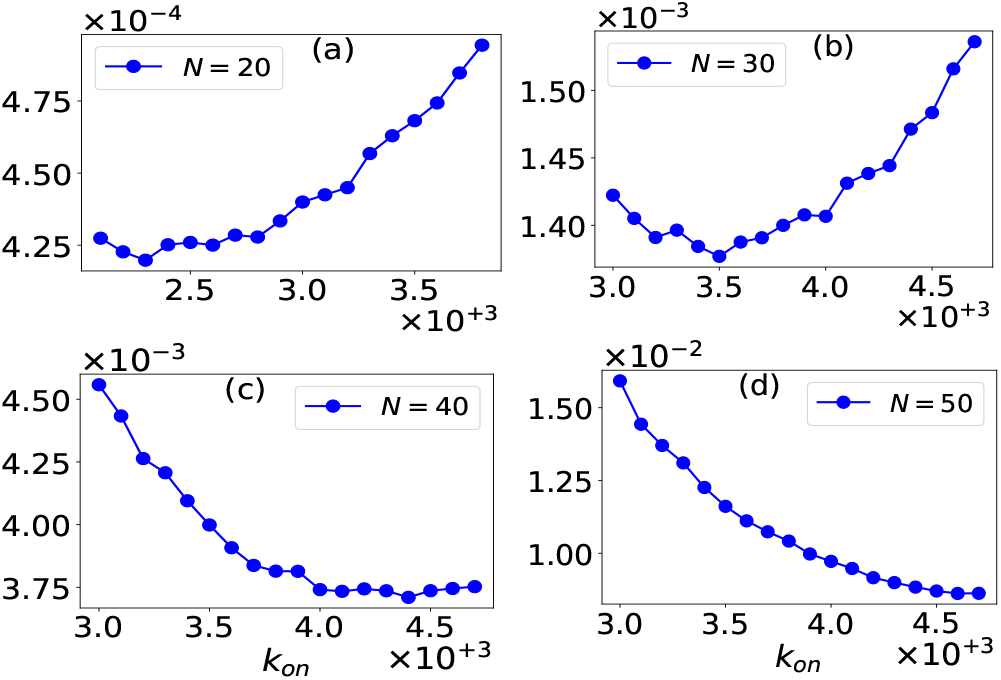
Non-monotonic behavior of the mean nucleation time ⟨*τ_s_*⟩ with increasing on-rate. The off-rate was decreased simultaneously from 52000 to 35000, linearly. The minimum shifts to the right as we increase the nucleus size (see legends).

However, we implement the same percentage of variation in our on/off rates. So, we varied the on-rate from 3000 to 4600 and off-rate from 52000 to 35000 (linearly) to get *τ_s_*. Each of the points on each curve is obtained by the averaging over 10^4^ nucleation events. As the on-rate increases (simultaneously decreasing the off-rate) we see a decrease in nucleation time at first, but then for higher values of on-rates (and correspondingly lower values of off-rates), we observe an increase in the mean sickling time (see fig.10.b,c).

This unexpected non-monotonic behavior is due to the low on-rate (and high off-rate) at low temperatures, resulting in slow polymerization, making the mean nucleation time longer for lower temperatures. Whereas, at a higher temperature, the on-rate being high (and off-rate being low), the monomers quickly polymerize to form the critical nucleus. At a further increase of temperature, the monomer pool is exhausted very fast by forming many short oligomers, and an accompanying low off-rate makes these oligomers very stable. As a result, the system gets trapped in a jammed state where no significant monomer addition/dissociation can occur. However, unless a monomer can dissociate from one oligomer and reassociate with another oligomer (an effective monomer exchange), the oligomers cannot grow further. This effect delays the nucleation time.

In between these two extreme cases of large ⟨*τ_s_*⟩, owing to two different mechanisms, occurs a minimum where the nucleation, time is short. As we increase the nucleus size (see fig.10 fig.11), the minimum shifts towards higher on-rate. This is because, for the stability of larger polymers (which is a precursor to the nucleus), a high on-rate and low off-rate are preferable. However, with a decrease in monomer number, the minimum shifts towards the left, i.e., towards lower on-rate (result not shown here). This is because the monomer pool is exhausted faster due to the formation of trapped states. Hence, for low monomer concentration, a slow polymerization (low on-rate) will be suitable for nucleus formation.

**FIG. 11.**
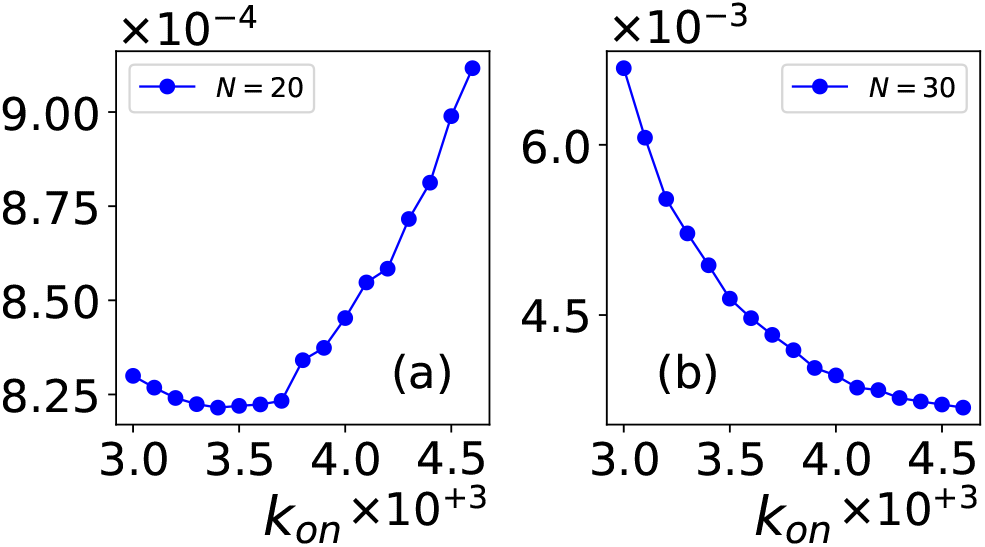
Non-monotonic behavior of the mean nucleation time ⟨*τ_s_*⟩ with increasing on-rate for the multi-filament case. The off-rate was decreased simultaneously from 52000 to 35000, linearly. The shifting of the minimum to the right is similar to that of the single-filament case.

Figure11 shows the mean first-assembly time for the case of 3-filament nucleus with *N* = 20 and 30. In these cases, the minimum shifts to the right and appears at a higher on-rate value than the corresponding mono-filament cases of the same *N* (compare fig.10b and fig.11b). This is because a larger number of stable oligomers must form to produce three oligomers of size *N* or larger. Note that here we defined the nucleation time as the earliest time when the system has three oligomers of size ≥ *N*. So, a relatively higher on-rate is suitable for stable oligomers than the mono-filament case. It should be noted that the non-monotonic dependency obtained in fig.10 and fig.11 are possible only if we take the onrate and off-rate to be temperature-dependent. For a constant off-rate, as claimed in ref.[5], one does not find any minimum in the mean sickling-time curve. However, a constant on-rate and a temperature-dependent off-rate can yield a similar minimum as in fig10, 11.

## DISCUSSION

Our KMC simulations, also supported by the BD model, show that large lag times could occur in two distinct scenarios : a) when monomer association rate *k_on_* is low and dissociation rate *k_off_* is high, and b) when *k_on_* is high and *k_off_* is low. In the first case, both primary nucleation and polymer growth rates are low, which is central to Szabo’s [17] model. This regime occurs when the temperature is low or when the initial monomer concentration is low. In practice, this can occur in microdroplet experiments or even inside the small volume of RBC. However, we focused on case b, which can occur when the temperature is high and as a result, *k_on_* is high and *k_off_* is low. In this regime, the system quickly reaches the jammed state leading to similar long-tailed skewed distribution. However, we also showed that the jamming effect gets neutralized if the initial monomer concentration is high.

It is interesting that the lag time distributions, obtained in our simulations, are very similar in these two regimes of high and low polymerization rates. Further, they also agree qualitatively with the trends observed in experiments [11, 25].

Note that the long tails in both the polymerization regimes (high and low) results from stochasticity of rare events. For the low polymerization rate, it is established [17] that it is the low copy number of nuclei, which leads to stochasticity. For rapid polymerization (and slow dissociation), it is the rare dissociation events that lead to stochasticity. In our model, we ignored the branching of polymers and the cooperative growth of the HbS filaments. Inclusion of these additional features are challenging, but we do not think these effects will change our results qualitatively, except making the overall polymerization process even faster.

The emergence of a minimum, although a shallow one, in our mean sickling time versus *k_on_* (equivalently temperature) plot is interesting, but it is difficult to speculate if this could be useful for therapeutic purposes. Treatments against RBC sickling aim to prevent the polymerization of HbS or delay its polymerization, i.e., extending the lag time. Ref[11] has proposed that the lag time can be extended by lowering HbS concentration via dilation of the RBC. Our simulations show that raising the temperature (and thereby the polymerization rate) can also extend the lag time, although it could be impractical to raise the patient’s body temperature artificially. Also, we have assumed that the observed in vitro poly-/depolymerization rates will be valid for in vivo situation also, which needs to be checked experimentally.

In general, experiments on temperature-dependent sickling can help us understand the pathophysiology of RBC sickling. While the increase of lag/sickling time at a low temperature has been reported before [12], how lag/sickling time behaves at a higher temperature, to the best of our knowledge, is not known and calls for new controlled experiments. In practice, patients with sickle cell disease are advised to avoid sudden exposure to cold or low temperatures, which triggers sickling. But how this cold stress induce sickling is unclear because naive generalization from in vitro experiments at low temperature would imply a delay of the lag times and hence slow sickling rate.

## ACKNOWLEDGEMENTS

AS and AB would like to thank Science and Engineering Research Board (SERB), India (Project No. MTR/2019/00135), and UGC-India, respectively, for financial support. We also acknowledge Space-Time-2 supercomputing facility at IIT Bombay, where most of the computations were carried out.

